# Infectome analysis of bat kidneys from Yunnan province, China, reveals close relatives of Hendra-Nipah viruses and prevalent bacterial and eukaryotic pathogens

**DOI:** 10.1101/2025.01.10.632301

**Authors:** Guopeng Kuang, Tian Yang, Weihong Yang, Jing Wang, Hong Pan, Yuanfei Pan, Qin-yu Gou, Wei-chen Wu, Juan Wang, Lifeng Yang, Xi Han, Yao-qing Chen, John-Sebastian Eden, Edward C. Holmes, Mang Shi, Yun Feng

## Abstract

Bats are natural reservoirs for a wide range of microorganisms, including many notable zoonotic pathogens. However, the infectome of bat kidneys remains poorly understood. To address this gap, we performed meta-transcriptomic sequencing on kidney tissues from 142 bats, spanning ten species sampled at five locations in Yunnan province, China. This analysis identified 22 viral species, including 20 novel viruses, two of which represented newly discovered henipaviruses closely related to the highly pathogenic Hendra and Nipah viruses. These henipaviruses were found in the kidneys of bats inhabiting an orchard near villages, raising concerns about potential fruit contamination via bat urine and transmission risks to livestock or humans. Additionally, we identified a novel protozoan parasite, tentatively named *Klossiella yunnanensis*, along with two highly abundant bacterial species, one of which is a newly discovered species—*Flavobacterium yunnanensis*. These findings broaden our understanding of the bat kidney infectome, underscore critical zoonotic threats, and highlight the need for comprehensive, full-spectrum microbial analyses of previously understudied organs to better assess spillover risks from bat populations.

## Introduction

Bats (order *Chiroptera*) are one of the most diverse and abundant groups of mammals, comprising nearly 1,500 species with a near global distribution^1^. Bats are also well-known natural reservoirs for a wide variety of microbial pathogens, a characteristic often attributed to their unique immune systems which maintain a delicate balance between host defenses and immune tolerance to viral infections^2–4^. Importantly, bats have been implicated in a number of major emerging disease outbreaks, including Hendra^5^, Nipah^6^, Marburg^7^ and Ebola^7^ virus disease, severe acute respiratory syndrome (SARS)^8^, Middle East respiratory syndrome (MERS)^8^, and coronavirus disease 2019 (COVID-19)^9^. Indeed, comparative studies indicate that bats harbor a greater diversity of viruses than many other mammalian groups, underscoring their significance for zoonotic disease surveillance^10^.

Metagenomic approaches have greatly advanced the characterization of bat viromes, deepening our understanding of the diversity of bat-borne pathogens and their potential role in disease emergence and transmission^11–14^. As of October 2024, viral sequences from at least 31 families have been identified in 340 bat species across 111 countries^15^. Bat-borne viruses are transmitted to humans either through direct contact with bats or via so-called “intermediate” hosts, often linked to the ingestion of food or water contaminated with bat saliva, feces, or urine^16^. While much of this research has focused on the bat gut virome, viruses present in other body sites—such as the kidneys—also pose transmission risks.

Indeed, zoonotic viruses have been detected in bat kidneys and urine, including henipaviruses^17–21^, pararubulaviruses^21–26^, and betacoronaviruses^26^. As these kidney-associated pathogens can be excreted through urine they are at heightening risk of human exposure.

Beyond viruses, bats harbor a diverse array of bacteria, fungi, and protozoan parasites that infect bats or even humans^27,28^. A notable example is the psychrophilic fungus *Pseudogymnoascus destructans*, which has caused a devastating disease in bats and led to the deaths of millions of animals across eastern North America^28^. Although this fungus is not known to pose a direct threat to humans, disturbances during bat hibernation—such as flying during the day and gathering near cave and mine entrances in winter—may increase human-bat encounters. Additionally, zoonotic bacteria and protozoan parasites, such as members of *Leptospira*^29,30^ and *Toxoplasma*^31^, have been identified in bat kidneys. However, as with viruses, research on the bacteria and eukaryotic pathogens in bat kidneys remain sparse, highlighting a critical gap in our understanding of the diversity of bat pathogens.

Yunnan province, located in southwestern China and bordering a number of Southeast Asian countries, is renowned as a hotspot for bat diversity and bat-borne viral pathogens, including close relatives of Marburg virus^32^, SARS-CoV^33,34^, and SARS-CoV-2^9,35,36^. Herein, we utilized a meta-transcriptomics approach to investigate the total infectome—comprising viruses, bacteria, and eukaryotic pathogens—in bat kidneys collected from this geographic region. We further identified and characterized potential human pathogens of notable zoonotic risk and explored interactions between viruses and their protozoan parasite hosts, offering valuable insights into the complexity of the bat kidney infectome.

## Results

### Bat species identification

Between 2017 and 2021, kidney tissues were sampled from 142 individual bats across five cities/counties in Yunnan province, China (Fig. 1a and Supplementary Table 1). Species identification was initially performed by recovering partial cytochrome c oxidase I (*COX1)* gene sequences using targeted PCR assay and Sanger sequencing. Phylogenetic analysis of full-length *COX1* sequences, generated from meta-transcriptomic sequencing, confirmed the presence of ten bat species spanning five genera and three families (Fig. 1b and Supplementary Table 1). Based on mitochondrial sequences and sampling locations, the samples were pooled into 20 groups for sequencing library constructions, with each group containing 2 to 8 individuals (Supplementary Table 1). Meta-transcriptomic sequencing of total RNA extracted from these pools yielded an average of 56.33 million clean non-rRNA reads, totaling approximately 1.13 billion clean non-rRNA reads.

**Fig. 1.**
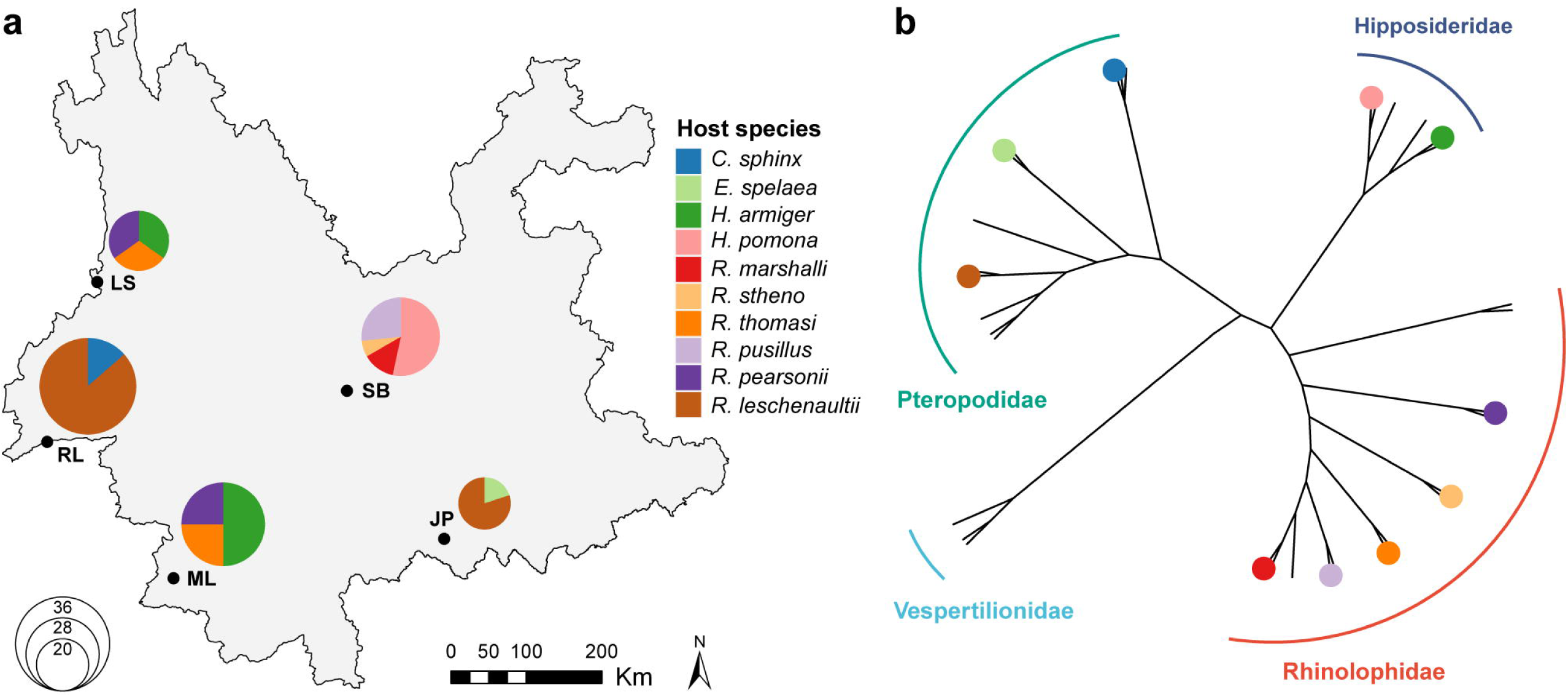
Bat kidney sampling and species identification. **(a)** Map showing the five sampling locations in Yunnan province, China, with pie charts indicating the species composition of the bats sampled at each site. The basemap shapefile used in ArcGIS was obtained from the publicly available GADM data set (https://gadm.org/download_country.html). **(b)** Unrooted phylogenetic tree inferred from full-length *COX1* gene sequences of bat kidney samples analyzed in this study. Colors correspond to different bat species, matching the color scheme used in the pie charts. Branch lengths are scaled to the number of nucleotide substitutions per site.

### Overview of the Yunnan bat kidney infectome

Meta-transcriptomic analysis of the bat kidneys identified a diverse microbial community (Fig. 2 and Supplementary Table 2). Based on our detection criteria (see Methods), microbes were detected in 18 of the 20 analyzed libraries, comprising 0.06% to 1.28% of total clean non-rRNA reads per library (Fig. 2a). Two libraries—one from *Hipposideros armiger* (8 individuals, LS) and another from *Rhinolophus stheno* (2 individuals, SB)—showed no microbial presence. RNA viruses dominated the microbial community, with 20 species from 12 families identified, as well as one DNA virus, one reverse transcribing virus, two bacterial species, and one eukaryotic species (Fig. 2b). Among these, the eukaryote genus *Phyllobacterium* was the most frequently detected, present in 13 (65%) libraries. Notably, one library contains a total of 12 microbial species, all of which were viruses (Fig. 2c).

**Fig. 2.**
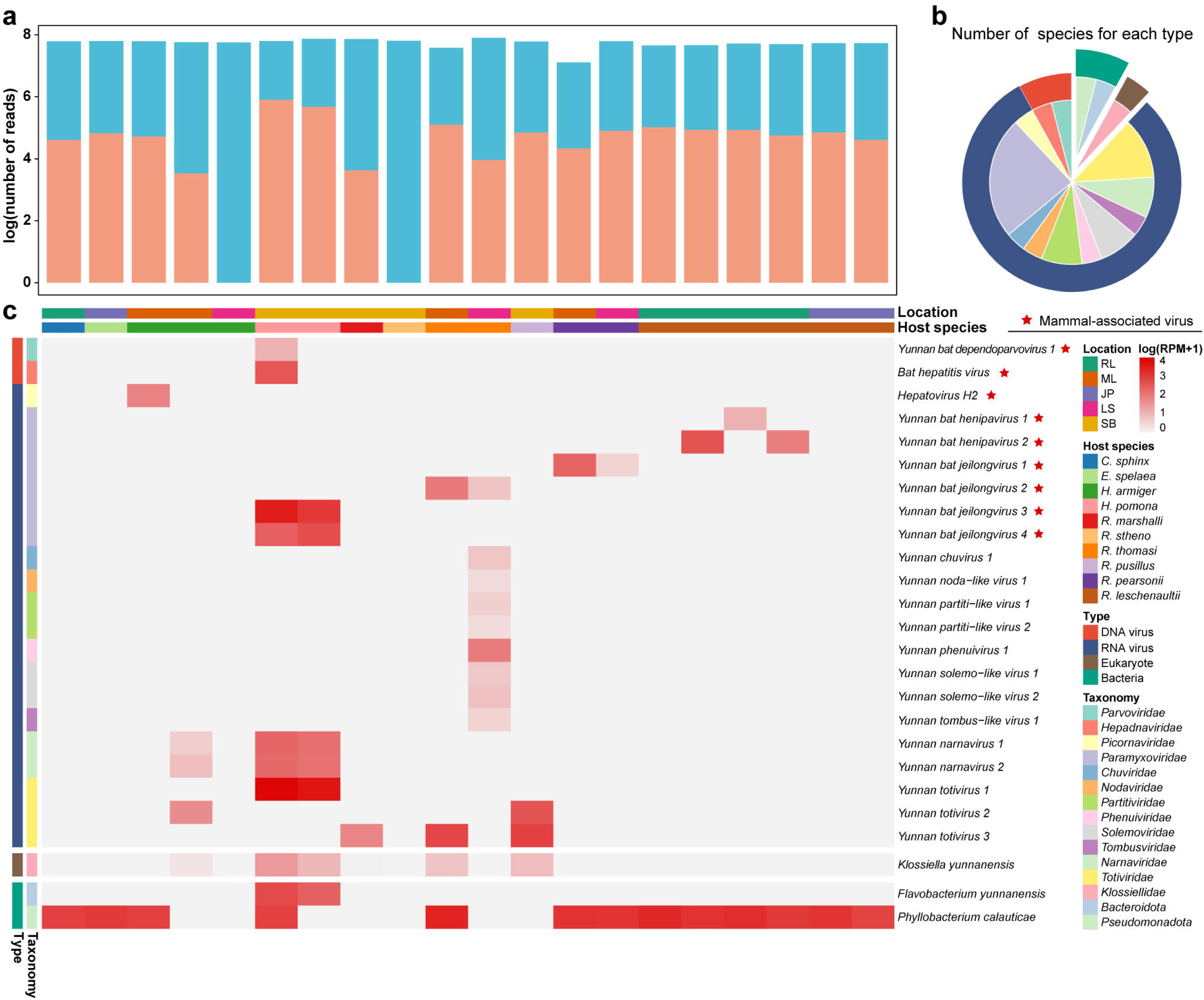
Overview of the bat kidney infectome. **(a)** Numbers of total reads (light blue) and microbial reads (orange) for each library. **(b)** Count of viral, bacterial, and eukaryotic microbial species detected. **(c)** Heatmap illustrating the distribution and relative abundance of viral, bacterial, and eukaryotic microbes, represented as RNA abundance (RPM: reads per million non-rRNA reads) in each library. Host species and orders are labeled at the top and color-coded according to their respective categories.

### Virome of bat kidneys

We identified 22 viral species across 12 families in bat kidneys (Fig. 2 and 3). These included six RNA viruses from the *Paramyxoviridae*, three from the *Totiviridae*, two each from the *Partitiviridae, Solemoviridae*, and *Narnaviridae*, and one each from the *Phenuiviridae, Chuviridae, Nodaviridae, Picornaviridae*, and *Tombusviridae*. Additionally, we discovered one DNA virus from the *Parvoviridae* and one reverse transcribing virus from the *Hepadnaviridae* (Fig. 2, 3). Of these, 20 species (90.91%) spanning 10 families were newly identified per ICTV (International Committee on Virus Taxonomy) species demarcation criteria (Supplementary Table 2).

**Fig. 3.**
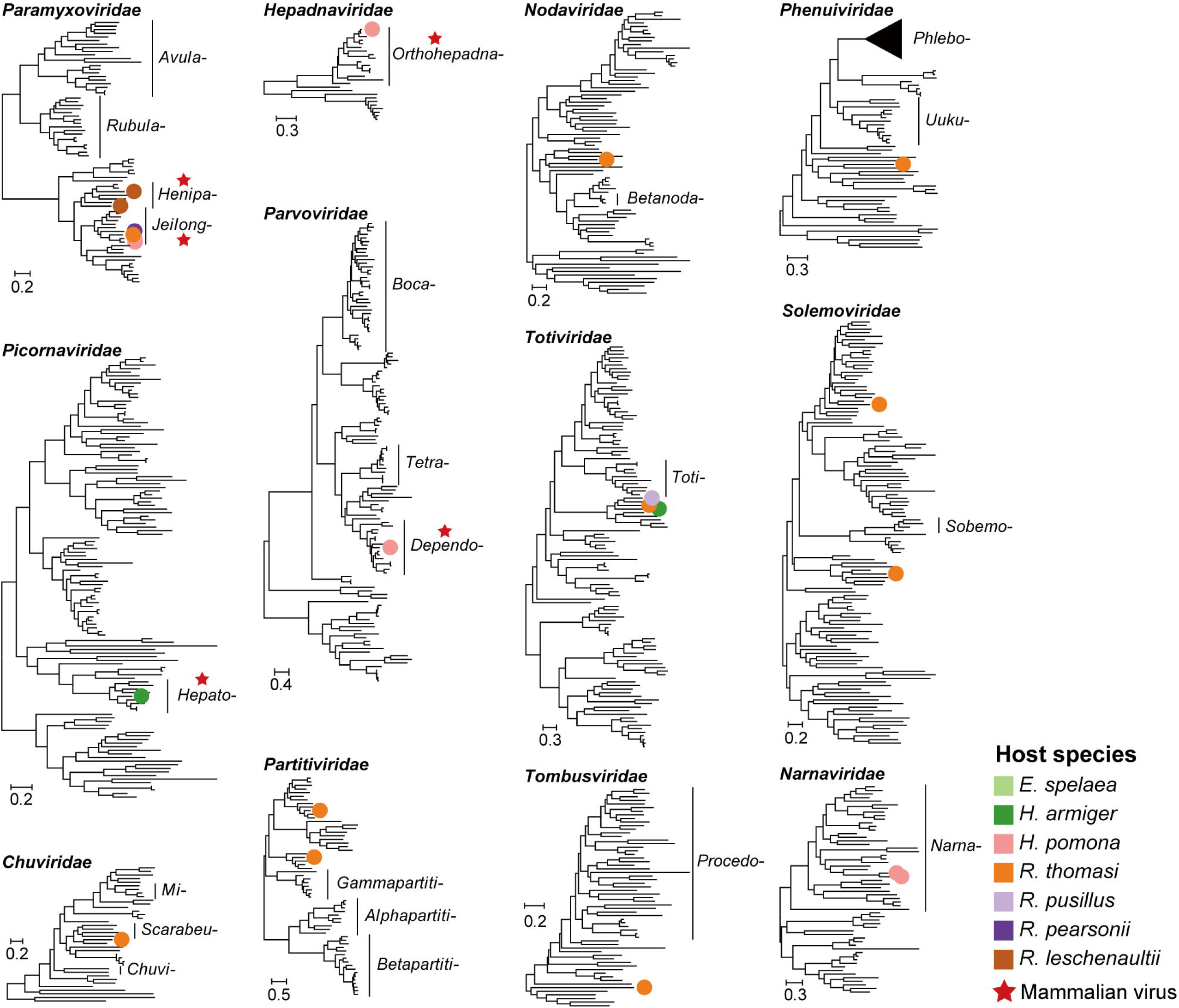
Phylogenetic diversity of viruses identified in this study. Phylogenetic trees of viruses from 12 virus families estimated using the maximum likelihood method based on conserved protein sequences (RdRp for RNA viruses, NS1 for *Parvoviridae*, and DNA polymerase for *Hepadnaviridae*). Colored dots on the trees, corresponding to host genera as indicated in the legend, represent viral species identified in this study. Red stars mark members of known mammal-associated viral lineages. All trees are mid-pointed rooted for clarity only with horizontal branch lengths depicting the number of amino acid substitutions per site.

Phylogenetic analyses revealed that nine species (40.91%) were related to known mammal-associated viruses, representing one reverse transcribing virus, one DNA virus and seven RNA viruses (Fig. 3). The *Paramyxoviridae* exhibited the highest diversity, with two species from the genus *Henipavirus* and four from the genus *Jeilongvirus* identified. Notably, the two newly identified henipaviruses showed relatively close evolutionary links to two human pathogens—Hendra virus (HeV, 52.23–56.94% identity in the L protein) and Nipah virus (NiV, 52.17–57.03%) (Fig. 3). Interestingly, we identified a hepatotropic virus, denoted Bat hepatitis virus variant YNBS16, in bat kidneys. Phylogenetic analysis revealed that this sequence was closely related to a sequence, ZYPR16 (Clade BtHBV 7), previously identified in bat livers (Extended Data Fig. 1).

There was considerable variation in diversity among viral families (Fig. 2b, c and 3). While the *Paramyxoviridae* dominated the sampled obtained, in many cases the highest abundance members of this family were not associated with the infection of vertebrates, including Yunnan narnavirus 1 and 2 (*Narnaviridae*) and Yunnan totivirus 1–4 (*Totiviridae*). Yunnan totivirus 1 was especially abundant in pools YNBS16 (RPM = 7393.98) and YNBS17 (RPM = 4171.90) (Fig. 2c), indicating that its presence was unlikely due to environmental contamination or dietary origin.

### Characterization of newly identified henipaviruses

Among the viruses identified, we focused on those with potential emergence risks based on their phylogenetic relationship to known high-impact human pathogens, specifically Yunnan bat henipavirus 1 and 2. Of the 20 pooled libraries, one (YNBS03) was positive for Yunnan bat henipavirus 1, while two (YNBS02 and YNBS04) contained reads corresponding to Yunnan bat henipavirus 2. These positive pools were all derived from the kidneys of *Rousettus leschenaultii* bats inhabiting an orchard near villages in RL (WD) (Fig. 2c).

Using henipavirus genome sequences assembled from these libraries, primers were designed to further examine individual kidneys through qRT-PCR. The results revealed that one kidney from pool YNBS03 (sample WDBS1745), one from YNBS02 (sample WDBS1733), and two from pool YNBS04 (samples WDBS1762 and WDBS1769) tested positive for henipavirus. Further testing of other organs (heart, liver, lung, gut, and brain) from the same individuals (WDBS1733 and WDBS1745) using qRT-PCR and meta-transcriptomic sequencing confirmed the multi-organ presence of henipaviruses within these bats, with the exception of brain tissues (Table 1). Notably, the kidneys exhibited significantly higher viral abundance compared to other organs, suggesting that they are the primary site of henipavirus replication within the host.

**Table 1.** Detection of henipavirus in various organs within individual bats.

| Library | Organ | Virus query | Length<br>(bp) | Number<br>of reads | qRT-PCR<br>(Ct) | Nested-PCR |
| --- | --- | --- | --- | --- | --- | --- |
| WDBN1745 | brain | Yunnan bat henipavirus 1 | 19755 | 0 | NoCt | - |
| WDBX1745 | heart | Yunnan bat henipavirus 1 | 19755 | 0 | NoCt | - |
| WDBG1745 | liver | Yunnan bat henipavirus 1 | 19755 | 0 | NoCt | - |
| WDBF1745 | lung | Yunnan bat henipavirus 1 | 19755 | 16 | 33.35 | - |
| WDBC1745 | gut | Yunnan bat henipavirus 1 | 19755 | 0 | NoCt | - |
| WDBS1745 | kidney | Yunnan bat henipavirus 1 | 19755 | 3953 | 23.82 | + |
| WDBN1733 | brain | Yunnan bat henipavirus 2 | 17723 | N/A | NoCt | - |
| WDBX1733 | heart | Yunnan bat henipavirus 2 | 17723 | 16 | 31.37 | + |
| WDBG1733 | liver | Yunnan bat henipavirus 2 | 17723 | 15 | 26.04 | + |
| WDBF1733 | lung | Yunnan bat henipavirus 2 | 17723 | 20 | 30.34 | + |
| WDBC1733 | gut | Yunnan bat henipavirus 2 | 17723 | 29 | 30.74 | + |
| WDBS1733 | kidney | Yunnan bat henipavirus 2 | 17723 | 161657 | 16.91 | + |

The complete genomes of Yunnan bat henipavirus 1 and 2 were successfully assembled from individual kidney samples WDBS1745 and WDBS1733, achieving mean sequencing depths of 27.99 fold and 1,274.77 fold, respectively (Fig. 4a). These two sequences were designated as Yunnan bat henipavirus 1 variant WDBS1745 and Yunnan bat henipavirus 2 variant WDBS1733. The open reading frames (ORFs) and gene arrangements of both viruses were consistent with other members of the genus *Henipavirus*, with each encoding six proteins (Fig. 4a).

**Fig. 4.**
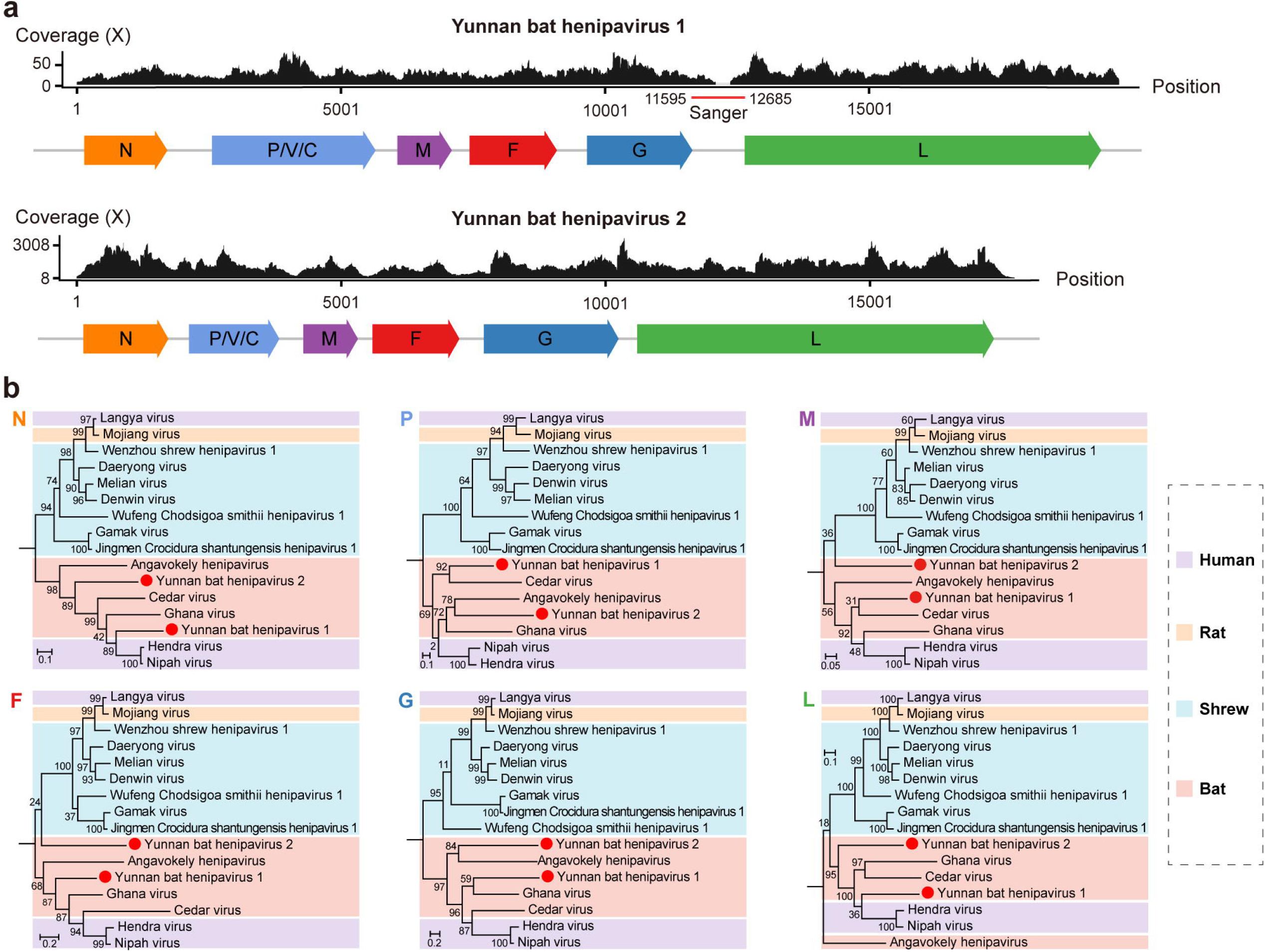
Characterization of the novel henipavirus species examined in this study. (a) Genome organization and sequencing coverage of two novel henipavirus species. Coverage across the full-length genome is displayed, with open reading frames (ORFs) depicted as colored arrows below the coverage plots. Regions confirmed by Sanger sequencing for Yunnan bat henipavirus 1 are marked with a red bar beneath the coverage graph. (b) Maximum likelihood phylogenetic trees estimated using amino acid sequences of each gene within the genus *Henipavirus*, rooted with J-virus (not shown). Color blocks indicate different species groups, and newly identified viruses are marked with solid red circles. All trees are mid-pointed rooted for clarity only with horizontal branch lengths depicting the number of amino acid substitutions per site.

Phylogenetic analysis of all six genes revealed a clear separation between the predominantly rodent-associated and bat/human-associated clades of the genus *Henipavirus* (Fig. 4b). Notably, the newly identified viruses formed distinct lineage, generally grouping with other bat-hosted henipaviruses, including the zoonotic pathogens HeV and NiV, both known for their high mortality rates in humans^37^. Yunnan bat henipavirus 1 was most closely related to HeV and NiV in the N (70.33–71.33% amino acid identity) and L proteins (56.94–57.03% amino acid identity), which underscores its potential risk as an emerging pathogen (Fig. 4b). However, the phylogenetic positions of Yunnan bat henipavirus 1 and 2 showed marked variability. In particular, Yunnan bat henipavirus 1 was most closely related to HeV and NiV in the N and L proteins trees, but occupied variable positions in the other trees. Although the bootstrap support for these groupings was generally weak, the topological movement of Yunnan bat henipavirus 1 among the henipaviruses likely reflects the action of recombination. Conversely, the phylogenetic positions of Yunnan bat henipavirus 2 was more consistent across gene trees and also exhibited a greater divergence from other bat henipaviruses.

### Identification and characterization of bacterial pathogens

Our meta-transcriptomic analysis also identified two bacterial taxa with relatively high abundance: *Flavobacterium* and *Phyllobacterium*. Phylogenetic analysis revealed that the *Flavobacterium* species forms a distinct branch closely related to *Flavobacterium ammonificans* (94.12% nucleic acid identity in the *ropB* gene) (Fig. 5a). This bacterium was tentatively classified as a novel species and named *Flavobacterium yunnanensis*. Similarly, the *Phyllobacterium* species was confirmed as *Phyllobacterium calauticae* based on 97.30% nucleic acid identity and phylogenetic placement of the *groEL* gene (Fig. 5b). Further transcriptomic profiling across all 20 pools demonstrated diverse gene expression patterns for these bacteria (Fig. 5c), indicating that they are metabolically active within the bat hosts.

**Fig. 5.**
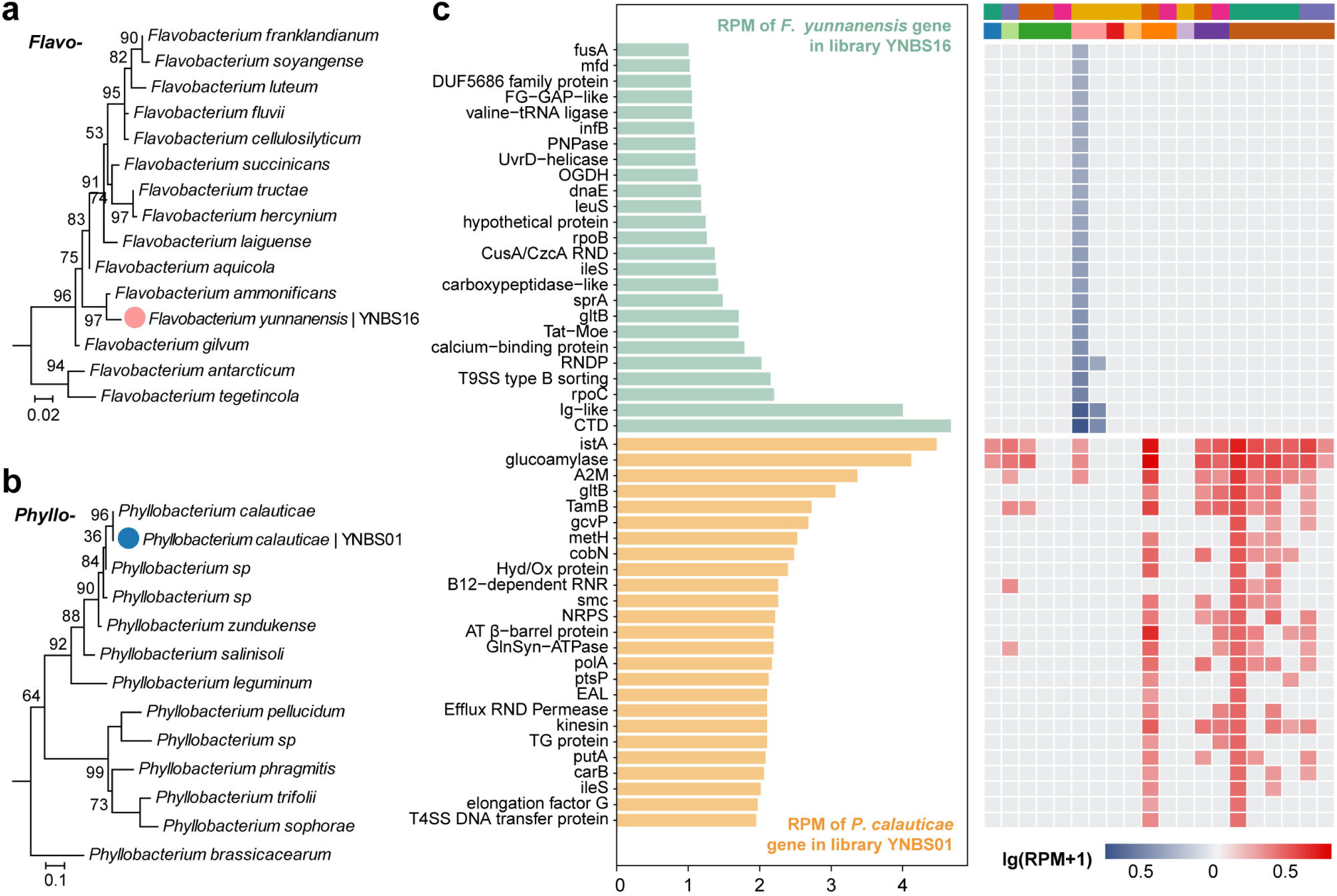
Gene expression profiles, prevalence and identification of the two bacterial microbes. (a) Maximum likelihood phylogenetic tree of the genus *Flavobacterium*, constructed using the rpoB gene. (b) Maximum likelihood phylogenetic tree of the genus *Phyllobacterium*, constructed using the groEL gene. (c) Top 25 expressed genes (measured in RPM) for *Flavobacterium yunnanensis* and *Phyllobacterium calauticae* in pools YNBS16 and YNBS01, respectively (left panel), compared with their expression in other positive pools (right panel).

### Eukaryotic microbe identified in bat kidneys

Analysis of genes *COX1* and cytochrome b (*cytB*) identified a protozoa pathogen closely related to the *Klossiella equi* of the family Klossiellidae (phylum Apicomplexa), known to infect the kidney of horses^38^. Phylogenetic and sequence divergence analyses revealed 87.7% nucleotide identity to *K. equi* (MH203050.1) in the *COX1* gene and 91.4% identity in the *cytB* gene. Based on these findings, the newly identified protozoan was proposed as a novel species, tentatively named *Klossiella yunnanensis* (Fig. 6a, b). *Klossiella* mitochondrial reads were detected in six pools, exhibiting uneven gene expression levels across different libraries (Fig. 6c).

**Fig. 6.**
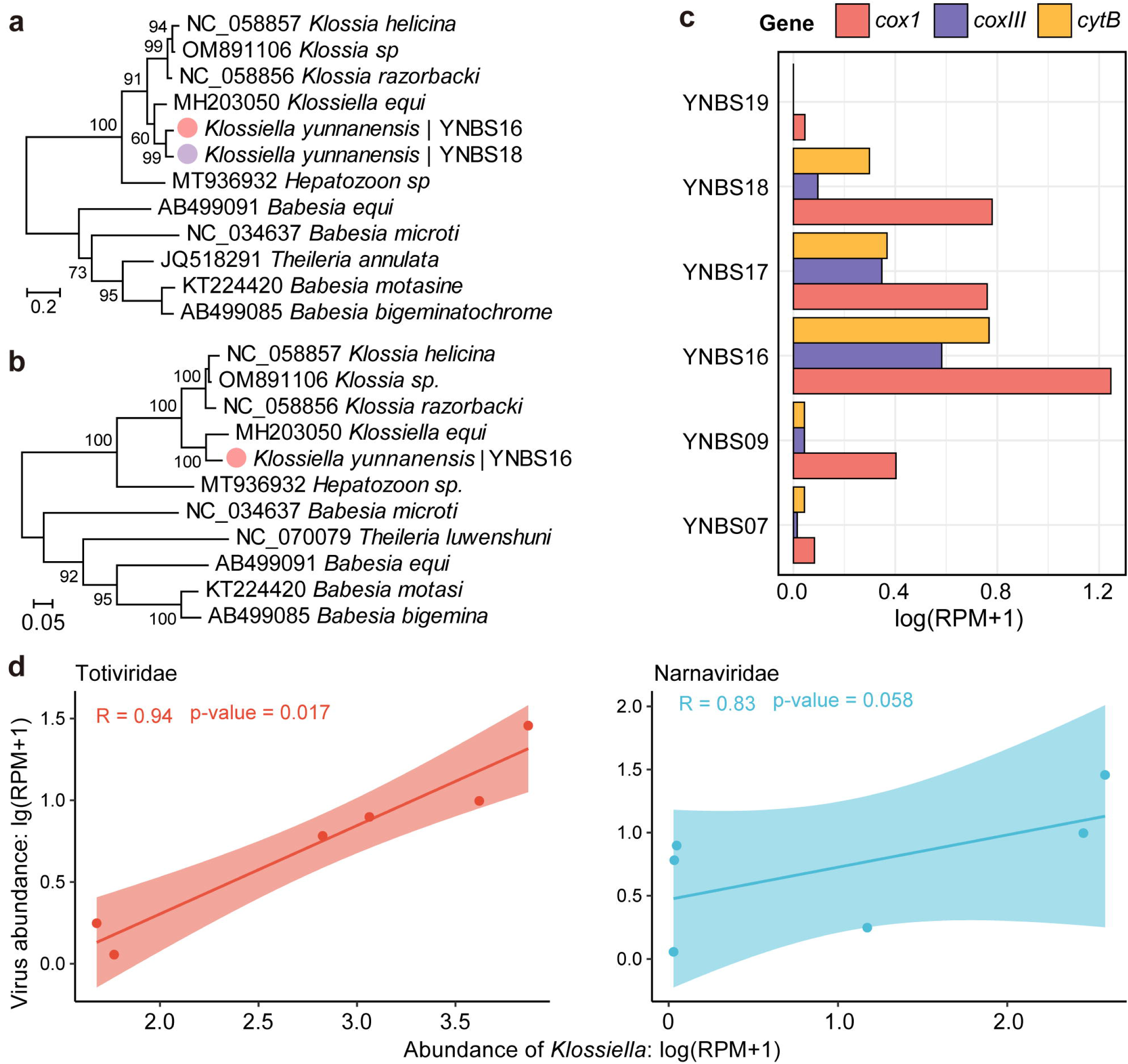
Identification and characterization of a eukaryotic microbe. (a, b) Phylogenetic trees of *Klossiella*, estimated using nucleotide sequences of the cox1 gene (a) and cytb gene (b). Colored dots indicate newly identified eukaryotic species, with colors corresponding to host genera. (c) Transcriptomic profiles of the *Klossiella* mitochondrion, represented as RPM, across positive pools. (d) Spearman’s correlation analysis showing the relationship between the total relative abundance (RPM) of totiviruses, narnaviruses, and *Klossiella yunnanensis*.

Interestingly, members of the viral families *Totiviridae* and *Narnaviridae*, known to naturally infect protozoa or fungi, showed co-occurrence with *K. yunnanensis* (Fig. 2c). To explore this relationship, we analyzed the correlation between the relative abundances (in RPM) of these viruses and *Klossiella* across the six positive pools. Strong positive correlations were observed, with Spearman’s ρ ranging from 0.83 to 0.94 (*p* < 0.05) (Fig. 6d), suggesting that the viruses in these families are likely hosted by *K. yunnanensis* rather than by bats.

## Discussion

There have been many studies of the presence of viruses, bacteria, and eukaryotic microbes (i.e., fungi and protozoan parasites) in various bat tissues, including the brain, lung, liver, rectum, feces, urine, throat, and fecal swabs^9,32–36,39,40^. In contrast, the kidneys have received comparatively little attention. Our meta-transcriptomic sequencing of bat kidneys revealed a diverse array of microorganisms, shedding light on the broader bat infectome. Although viruses were the predominant microbial group identified, only 9 of the 22 detected viral species were categorized as mammalian viruses. Notably, the mammal-associated viruses identified in the kidneys differed from those identified in the rectal tissues of the same individual bats^36^. These findings align with previous research showing that viruses from different families exhibit marked variation in their organ-specific distribution in bats^14^. As a consequence, these results underscore the importance of adopting a multi-organ approach to comprehensively understand the microbial diversity harbored by bats, particularly for identifying host-microbe interactions.

Of particular note, our study identified close relatives of Hendra and Nipah viruses colonizing bat kidneys. Nipah virus (NiV) are lethal pathogens that cause severe diseases in humans, including acute respiratory distress and encephalitis, with a mortality rate of 35-75%^37,41^. Similarly, Hendra virus (HeV) has caused multiple fatal outbreaks in humans and horses, including the death of veterinarians^37^. These viruses are naturally hosted by fruit bats (*Pteropus* species) and are typically transmitted to humans through bat urine or saliva, often via contamination of food sources^42,43^. HeV and NiV were first identified in Australia and Malaysia, respectively, and associated with Pteropus and other bat species^5,6^. Herein, we identified two related henipaviruses in *Rousettus leschenaultii* bats, marking the first detection of henipavirus genomes in bats from China. Previously, antibodies to Nipah or Nipah-like viruses have been reported in bats from multiple regions in China, including Yunnan, Guangdong, Hainan and Hubei provinces, suggesting potential exposure to such viruses^44^. Until now, however, no henipavirus genome sequences had been documented in bats from China. Notably, more distantly related viruses have been discovered in rodents and shrews, including Mojiang virus^45^ and Langya virus^46^, with the latter confirmed to infect humans. These findings highlight the significance of continued surveillance and genomic characterization of henipaviruses in bats, which are critical for understanding their potential spillover risk.

We also identified at least one bacterial species prevalent in bat kidneys. While the gut microbiota of bats has been extensively studied, less attention has been given to those of other organs, including the kidneys^16^. Previous research identified *Leptospira spp.* in bat kidney, supporting the hypothesis that bat kidneys may serve as a reservoir for zoonotic *Leptospira*^47–50^, and we previously detected pathogenic *Leptospira* in bat kidneys using nested PCR in individual tissue samples^51^. However, no *Leptospira*-associated reads were detected in the meta-transcriptomic sequencing of this study, possibly due to sample pooling which might obscure the detection of low-abundance microbes. Instead, we identified *Flavobacterium* and *Phyllobacterium*, of which *Phyllobacterium calauticae* exhibited relatively high abundance and prevalence (Fig. 2 and 5). *Phyllobacterium calauticae* is an aerobic, motile bacterium isolated from microaerophilic freshwater sediments, adapted to efficiently utilize oxygen in low-oxygen environments^52^. This is not unprecedented, as *Listeria monocytogenes*, an environmentally ubiquitous bacterium, has previously been isolated from various wild animals, including bat kidneys^53,54^.

Previous studies have shown that bats harbor a diverse range of protozoan parasites, some of which are capable of infecting humans^55^. However, research on protozoan parasites present in bat kidneys is limited. *Toxoplasma gondii*, a zoonotic protozoan parasite, has been detected in bats collected in Yunnan^56^, and herein we identified a protozoan parasite, tentatively named *Klossiella yunnanensis*, in six (30%) of the libraries. Phylogenetic analysis suggests that *K. yunnanensis* is closely related to species known to infect horses, *K. equi*, which is generally considered non-pathogenic but can cause kidney alterations in cases of heavy infection^38^. The pathogenicity of this eukaryotic parasites to humans or even bats remains unclear.

In addition, viruses from the families *Totiviridae* and *Narnaviridae*^57,58^, which are known to infect a wide range of non-vertebrate host types including protozoan parasites, were detected in high abundance in bat kidneys. Spearman’s correlation analysis of relative abundances indicated that these viruses were associated with *K. yunnanensis* rather than the bat hosts themselves (Fig. 6d). This highlights the importance of conducting studies of the total infectome to better elucidate the interactions between viruses within an animal and their potential relationship with the primary host.

Our study has several limitations. Uneven sampling across locations and bat species—in which each species was sampled at only one or two sites—complicated our ability to assess virus distribution, compare viral compositions between species, and identify transmission networks. In addition, the analysis of pooled samples with varying numbers of individuals per pool prevented us from determining whether the detected microbes were co-infections within individual bats or from different individuals. The unequal pool sizes may have also influenced the accuracy of microbial quantification. Finally, the lack of a complete genome assembly for the newly discovered eukaryotic parasite, coupled with the absence of a reference genome in existing databases, limited our ability to accurately quantify its abundance. Despite these limitations, our study offers the first comprehensive characterization of the bat kidney infectome, providing a foundation for more effective discovery and characterization of potential bat-borne pathogens.

## Materials and methods

### Ethics statement

This research, including the specimen collection and processing procedures, was reviewed and approved by the Ethics Committee of the Yunnan Institute of Endemic Disease Control and Prevention (File No. 20160002). All experiments were conducted with the approval of the Biosafety Committee of the same institute.

### Sample collection

Five sampling sites in Yunnan province were selected, denoted RL, ML, SB, LS, and JP (Fig. 1a). From 2017 to 2021, sticky nets were deployed around orchards and caves to capture bats, which were promptly removed after being trapped. Initial identification of bat species was performed by experienced field biologists based on morphological characteristics. Captured bats were then transported to the laboratory, euthanized by intracardiac delivery of sodium pentobarbitone, and dissected. Kidney tissues were collected and stored at −80LJ°C until further analysis. Preliminary species identification was confirmed by sequencing the cytochrome c oxidase I (*COX1*) gene for each specimen^59^. Mammalian species confirmation was achieved using *de novo* assembled *COX1* gene contigs. The final clean cox1 contigs were compared against the database within the BARCODE OF LIFE DATA SYSTEM (BOLDSYSTEMS)^60^, and phylogenetic analyses were conducted using PHYML 3.0^61^ for species identification.

### Meta Dtranscriptomic sequencing

Individual tissues were initially organized into sample groups based on species identification and collection location. Specifically, 142 kidney tissues were grouped into 20 libraries, each comprising 2 to 8 individuals (Supplementary Table 1). Total RNA was extracted and purified from each pool using the RNeasy Plus Universal Mini Kit (Qiagen, Germany). RNA libraries were constructed using the Zymo-Seq RiboFree™ Total RNA Library Kit (No. R3003) (Zymo Research, USA), following the manufacturer’s instructions. These libraries were sequenced using paired-end 150 bp reads on the Illumina NovaSeq 6000 sequencing platform.

### Characterization of total infectomes

Adapter sequences were removed from the sequencing reads, and initial quality control was performed using the pipeline implemented in bbduk.sh (https://sourceforge.net/projects/bbmap/). Duplicate reads were filtered out using cd-hit-dup with default settings^62^. rRNA reads were removed by mapping the processed reads against the SILVA rRNA database (Release 138.1) using Bowtie2 (version 2.3.5.1) in ‘--local’ mode^63^.

The remaining high-quality, non-rRNA reads were either (i) directly compared against the non-redundant protein (nr) database using DIAMOND Blastx^64^, or (ii) assembled into contigs using MEGAHIT (version 1.2.8)^65^ before comparison against the National Center for Biotechnology Information (NCBI) non-redundant protein (nr) database. An e-value threshold of 1×10^−5^ was set to maintain high sensitivity and minimize false positives.

For virus identification, contigs identified under the kingdom ‘Viruses’ were extracted, and those shorter than 600 bp were excluded to ensure the quality of virus genomes. The remaining overlapping contigs were merged into extended viral sequences using the SeqMan program implemented in the Lasergene software package version 7.1 (DNAstar, USA)^66^.

Each viral contig was classified at the species level based on the species demarcation criteria established by the ICTV for the respective viral genus^58^. For genera lacking explicit species demarcation criteria, a 90% amino acid identity threshold for the RdRP or replicase protein was applied (Supplementary Table 2). The abundance of these viral contigs was estimated by mapping reads back to the assembled genomes using Bowtie2 version 2.5.2 with ‘--end-to-end’ and ‘--very-fast’ settings. Alignments were sorted and indexed with SAMtools version 1.18 and visualized with Geneious Prime version 2020.2.4^67,68^.

For bacteria and eukaryotic microbes, we initially utilized MetaPhlAn version 4 to identify potential microbial taxonomy^69^. Complete reference genome sequences of the corresponding genera were subsequently downloaded from GenBank and used as templates for read mapping and gene abundance estimation with Bowtie2 (version 2.5.2)^63^. Highly conserved regions, such as rRNA genes, were excluded from the reference genome sequences before conducting mapping analyses. Based on the mapping results, we generated consensus sequences of marker genes (e.g., *rpoB*, *groEL*, *recA*, and *gyrB*), which were then subjected to BLASTn comparisons against the NCBI nucleotide (nt) database to determine microbial taxonomy at the species level.

### Evolutionary analyses

To determine the evolutionary relationships of the newly identified microbes, reference nucleotide/amino acid sequences for microbial taxa in question were downloaded from the NCBI GenBank Database. In all cases, sequences were then aligned using MAFFT^70^, with the 5’ and 3’ unaligned regions (when present) removed manually and ambiguously aligned sequences excluded using TrimAl version 1.5.0^71^. Phylogenetic trees on these data were then estimated using the maximum likelihood method implemented in PHYML 3.0, employing the GTR model of nucleotide substitution and SPR branch swapping^61^. Node support was estimated using an approximate likelihood ratio test using Shimodaira–Hasegawa-like procedures.

### Characterization of henipaviruses

To assess the prevalence of novel henipaviruses in bats and in different organs, real-time quantitative reverse transcription PCR (qRT-PCR) and nested RT-PCR were performed on all individual kidney samples. Specific primers were designed using the virus genome sequences obtained from libraries YNBS02 (Yunnan bat henipavirus 1) and YNBS03 (Yunnan bat henipavirus 2). To investigate viral distributions across various bat organs, PCR detection and individual meta-transcriptomics assays were performed on the brain, heart, liver, kidney, and gut sample of the positive bats (WD1733 and WD1745). However, the library construction for the brain sample from WDBN1733 failed.

As the full-length sequence of Yunnan bat henipaviruses 1 was not initially obtained, PCR assays and Sanger sequencing were employed to complete it. The final genome consensus sequences were confirmed by mapping the reads against draft genome sequences, and viral abundance was estimated based on the number of reads mapped to genome^63^. For each complete genomes, potential open reading frames (ORFs) and coding arrangements were predicted using ORFfinder (https://www.ncbi.nlm.nih.gov/orffinder/) and annotated by blastp program (https://blast.ncbi.nlm.nih.gov/Blast.cgi). Phylogenetic trees for each gene were estimated following the standard protocol described above.

## Acknowledgments

This study was funded by grants from the National Key R&D Program of China (2024YFC2607502). Y. F. was supported by Yunnan Revitalization Talent Support Program Top Physician Project (XDYC-MY-2022-0074). M.S. was supported by the National Natural Science Foundation of China (82341118, 32270160), Natural Science Foundation of Guangdong Province of China (2022A1515011854), Shenzhen Science and Technology Program (KQTD20200820145822023), Major Project of Guangzhou National Laboratory (GZNL2023A01001), Guangdong Province “Pearl River Talent Plan” Innovation, Entrepreneurship Team Project (2019ZT08Y464), and the Fund of Shenzhen Key Laboratory (ZDSYS20220606100803007). E.C.H. was supported by a National Health & Medical Research Council (NHMRC) Investigator grant (GNT2017197) and by AIR@InnoHK administered by the Innovation and Technology Commission, Hong Kong Special Administrative Region, China.

We wish to thank the local Centers for Disease Control and Prevention in six trapping sites for their assistance in specimen collection.

## Author Contributions

Conceptualization, G.-P.K., T.Y., M.S., and Y.F.; Methodology, G.-P.K., J.W., M.S., and Y.F.; Investigation, G.-P.K., T.Y., W.-H.Y., J.W., H.P., Y.-F.P., W.-C.W., Y.-Q.C., M.S., and Y.F.; Writing – Original Draft, G.-P.K., T.Y., M.S., and Y.F.; Writing – Review and Editing, All authors; Funding Acquisition, M.S., and Y.F.; Resources (sampling), G.-P.K., T.Y., W.-H.Y., H.P., J.W., X.H., L.-F.Y., and Y.F.; Resources (Computational), G.-P.K., T.Y., Y.-F.P., Q.-Y.G, Y.-Q.C., and M.S.; Supervision, J.-S.E., E.C.H., M.S., and Y.F..

## Competing interests

The authors have declared that no competing interests exist.

**Additional information**

**Extended data**

**Extended Data Fig. 1**

## Supplementary information

**Supplementary Table 1.** Information of sample group and RNA library in this study.

**Supplementary Table 2.** Viruses in bat kidneys identified in this study.

---

**Extended Data Fig. 1.**
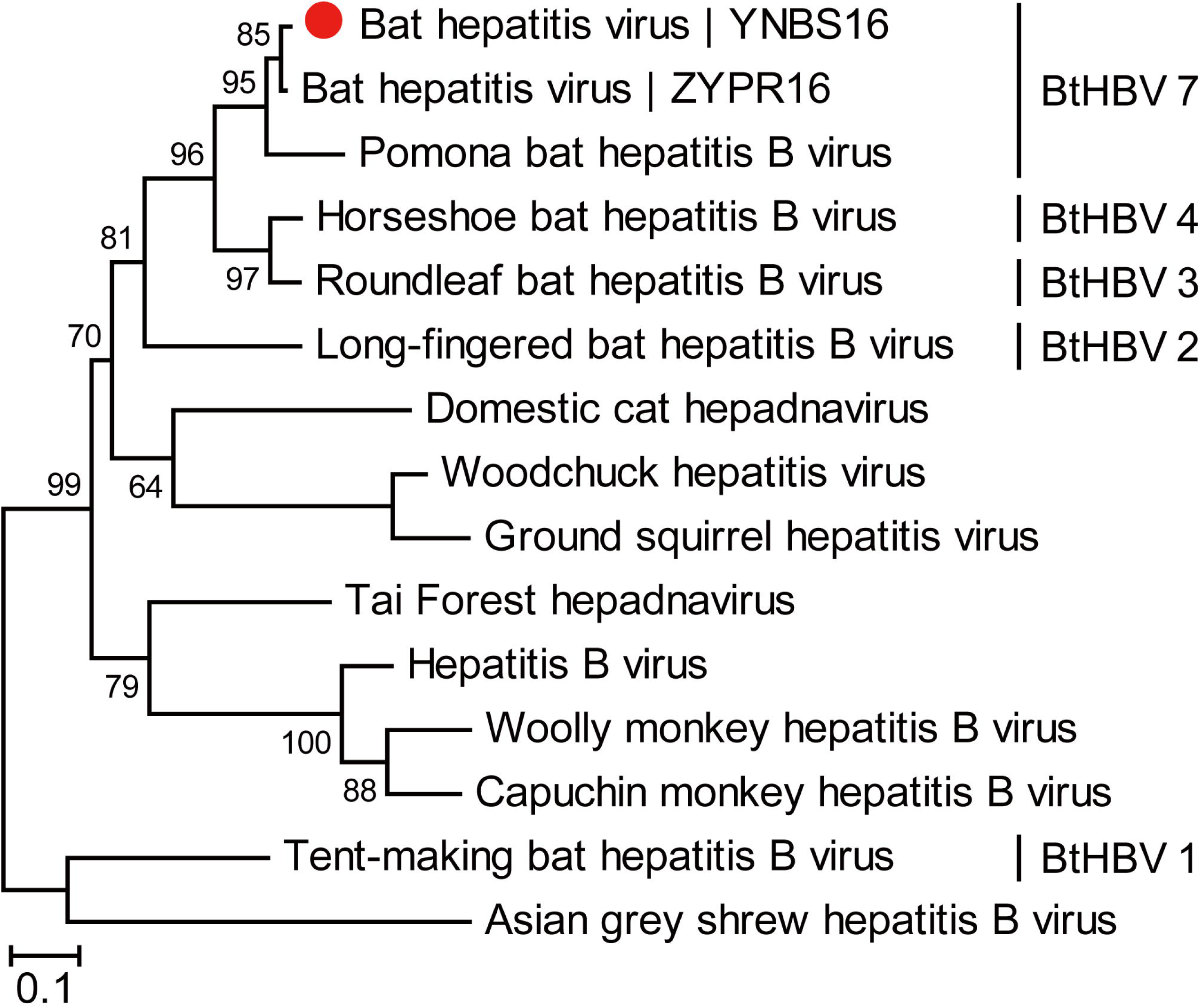
Maximum likelihood phylogenetic tree constructed based on amino acid sequences of the DNA polymerase within the genus *Orthohepadnavirus*. The newly identified virus in this study is marked with a solid red circle. Bat-derived strains and their corresponding clades—determined according to divergence limits used by ICTV as species demarcation criteria—are labeled on the right.

## References

1. Simmons, N.B. and A.L. Cirranello. 2024. Bat Species of the World: A taxonomic and geographic database. Version 1.6. https://zenodo.org/records/12802826

2. Calisher CH, Childs JE, Field HE, Holmes KV, Schountz T. Bats: important reservoir hosts of emerging viruses. Clin Microbiol Rev. 2006 Jul;19(3):531–45. doi: 10.1128/CMR.00017-06.

3. Hayman DT. Bats as Viral Reservoirs. Annu Rev Virol. 2016 Sep 29;3(1):77–99. doi: 10.1146/annurev-virology-110615-042203.

4. Irving AT, Ahn M, Goh G, Anderson DE, Wang LF. Lessons from the host defences of bats, a unique viral reservoir. Nature. 2021 Jan;589(7842):363-370. doi: 10.1038/s41586-020-03128-0. Epub 2021 Jan 20.

5. Halpin K, Young PL, Field HE, Mackenzie JS. Isolation of Hendra virus from pteropid bats: a natural reservoir of Hendra virus. J Gen Virol. 2000 Aug;81(Pt 8):1927–1932. doi: 10.1099/0022-1317-81-8-1927.

6. Chua KB, Koh CL, Hooi PS, Wee KF, Khong JH, Chua BH, Chan YP, Lim ME, Lam SK. Isolation of Nipah virus from Malaysian Island flying-foxes. Microbes Infect. 2002 Feb;4(2):145–51. doi: 10.1016/s1286-4579(01)01522-2.

7. Brainard J, Pond K, Hooper L, Edmunds K, Hunter P. Presence and Persistence of Ebola or Marburg Virus in Patients and Survivors: A Rapid Systematic Review. PLoS Negl Trop Dis. 2016 Feb 29;10(2):e0004475. doi: 10.1371/journal.pntd.0004475.

8. Cui J, Li F, Shi ZL. Origin and evolution of pathogenic coronaviruses. Nat Rev Microbiol. 2019 Mar;17(3):181–192. doi: 10.1038/s41579-018-0118-9.

9. Zhou P, Yang XL, Wang XG, Hu B, Zhang L, Zhang W, Si HR, Zhu Y, Li B, Huang CL, Chen HD, Chen J, Luo Y, Guo H, Jiang RD, Liu MQ, Chen Y, Shen XR, Wang X, Zheng XS, Zhao K, Chen QJ, Deng F, Liu LL, Yan B, Zhan FX, Wang YY, Xiao GF, Shi ZL. A pneumonia outbreak associated with a new coronavirus of probable bat origin. Nature. 2020 Mar;579(7798):270-273. doi: 10.1038/s41586-020-2012-7. Epub 2020 Feb 3. Erratum in: Nature. 2020 Dec;588(7836):E6. doi: 10.1038/s41586-020-2951-z.

10. Olival KJ, Hosseini PR, Zambrana-Torrelio C, Ross N, Bogich TL, Daszak P. Host and viral traits predict zoonotic spillover from mammals. Nature. 2017 Jun 29;546(7660):646-650. doi: 10.1038/nature22975. Epub 2017 Jun 21. Erratum in: Nature. 2017 Aug 31;548(7669):612. doi: 10.1038/nature23660.

11. Van Brussel K, Holmes EC. Zoonotic disease and virome diversity in bats. Curr Opin Virol. 2022 Feb;52:192–202. doi: 10.1016/j.coviro.2021.12.008. Epub 2021 Dec 23.

12. Wu Z, Yang L, Ren X, He G, Zhang J, Yang J, Qian Z, Dong J, Sun L, Zhu Y, Du J, Yang F, Zhang S, Jin Q. Deciphering the bat virome catalog to better understand the ecological diversity of bat viruses and the bat origin of emerging infectious diseases. ISME J. 2016 Mar;10(3):609–20. doi: 10.1038/ismej.2015.138. Epub 2015 Aug 11.

13. Wang J, Pan YF, Yang LF, Yang WH, Lv K, Luo CM, Wang J, Kuang GP, Wu WC, Gou QY, Xin GY, Li B, Luo HL, Chen S, Shu YL, Guo D, Gao ZH, Liang G, Li J, Chen YQ, Holmes EC, Feng Y, Shi M. Individual bat virome analysis reveals co-infection and spillover among bats and virus zoonotic potential. Nat Commun. 2023 Jul 10;14(1):4079. doi: 10.1038/s41467-023-39835-1.

14. Chen YM, Hu SJ, Lin XD, Tian JH, Lv JX, Wang MR, Luo XQ, Pei YY, Hu RX, Song ZG, Holmes EC, Zhang YZ. Host traits shape virome composition and virus transmission in wild small mammals. Cell. 2023 Oct 12;186(21):4662–4675.e12. doi: 10.1016/j.cell.2023.08.029.

15. Zhou S, Liu B, Han Y, Wang Y, Chen L, Wu Z, Yang J, 2022. ZOVER: the database of zoonotic and vector-borne viruses. Nucleic Acids Res. 50(D1):D943–D949. doi:10.1093/nar/gkab862

16. Han HJ, Wen HL, Zhou CM, Chen FF, Luo LM, Liu JW, Yu XJ. Bats as reservoirs of severe emerging infectious diseases. Virus Res. 2015 Jul 2;205:1–6. doi: 10.1016/j.virusres.2015.05.006.

17. Halpin K, Hyatt AD, Fogarty R, Middleton D, Bingham J, Epstein JH, Rahman SA, Hughes T, Smith C, Field HE, Daszak P; Henipavirus Ecology Research Group. Pteropid bats are confirmed as the reservoir hosts of henipaviruses: a comprehensive experimental study of virus transmission. Am J Trop Med Hyg. 2011 Nov;85(5):946–51. doi: 10.4269/ajtmh.2011.10-0567.

18. Marsh GA, de Jong C, Barr JA, Tachedjian M, Smith C, Middleton D, Yu M, Todd S, Foord AJ, Haring V, Payne J, Robinson R, Broz I, Crameri G, Field HE, Wang LF. Cedar virus: a novel Henipavirus isolated from Australian bats. PLoS Pathog. 2012;8(8):e1002836. doi: 10.1371/journal.ppat.1002836.

19. Barr J, Smith C, Smith I, de Jong C, Todd S, Melville D, Broos A, Crameri S, Haining J, Marsh G, Crameri G, Field H, Wang LF. Isolation of multiple novel paramyxoviruses from pteropid bat urine. J Gen Virol. 2015 Jan;96(Pt 1):24–29. doi: 10.1099/vir.0.068106-0.

20. Johnson RI, Tachedjian M, Rowe B, Clayton BA, Layton R, Bergfeld J, Wang LF, Marsh GA. Alston Virus, a Novel Paramyxovirus Isolated from Bats Causes Upper Respiratory Tract Infection in Experimentally Challenged Ferrets. Viruses. 2018 Nov 28;10(12):675. doi: 10.3390/v10120675. Erratum in: Viruses. 2021 Jun 23;13(7):1204. doi: 10.3390/v13071204.

21. Barr J, Smith C, Smith I, de Jong C, Todd S, Melville D, Broos A, Crameri S, Haining J, Marsh G, Crameri G, Field H, Wang LF. Isolation of multiple novel paramyxoviruses from pteropid bat urine. J Gen Virol. 2015 Jan;96(Pt 1):24–29. doi: 10.1099/vir.0.068106-0.

22. Barr JA, Smith C, Marsh GA, Field H, Wang LF. Evidence of bat origin for Menangle virus, a zoonotic paramyxovirus first isolated from diseased pigs. J Gen Virol. 2012 Dec;93(Pt 12):2590–2594. doi: 10.1099/vir.0.045385-0.

23. Baker KS, Todd S, Marsh GA, Crameri G, Barr J, Kamins AO, Peel AJ, Yu M, Hayman DT, Nadjm B, Mtove G, Amos B, Reyburn H, Nyarko E, Suu-Ire R, Murcia PR, Cunningham AA, Wood JL, Wang LF. Novel, potentially zoonotic paramyxoviruses from the African straw-colored fruit bat Eidolon helvum. J Virol. 2013 Feb;87(3):1348–58. doi: 10.1128/JVI.01202-12.

24. Baker KS, Tachedjian M, Barr J, Marsh GA, Todd S, Crameri G, Crameri S, Smith I, Holmes CEG, Suu-Ire R, Fernandez-Loras A, Cunningham AA, Wood JLN, Wang LF. Achimota Pararubulavirus 3: A New Bat-Derived Paramyxovirus of the Genus Pararubulavirus. Viruses. 2020 Oct 30;12(11):1236. doi: 10.3390/v12111236.

25. Federici L, Masulli M, De Laurenzi V, Allocati N. An overview of bats microbiota and its implication in transmissible diseases. Front Microbiol. 2022 Oct 20;13:1012189. doi: 10.3389/fmicb.2022.1012189.

26. Rizzo F, Edenborough KM, Toffoli R, Culasso P, Zoppi S, Dondo A, Robetto S, Rosati S, Lander A, Kurth A, Orusa R, Bertolotti L, Mandola ML. Coronavirus and paramyxovirus in bats from Northwest Italy. BMC Vet Res. 2017 Dec 22;13(1):396. doi: 10.1186/s12917-017-1307-x.

27. Dhivahar J, Parthasarathy A, Krishnan K, Kovi BS, Pandian GN. Bat-associated microbes: Opportunities and perils, an overview. Heliyon. 2023 Nov 18;9(12):e22351. doi: 10.1016/j.heliyon.2023.e22351.

28. US Fish and Wildlife Service. 17 January 2012. North American bat death toll exceeds 5.5 million from white-nose syndrome. US Fish & Wildlife Service news release. http://www.fws.gov

29. Bevans AI, Fitzpatrick DM, Stone DM, Butler BP, Smith MP, Cheetham S. Phylogenetic relationships and diversity of bat-associated Leptospira and the histopathological evaluation of these infections in bats from Grenada, West Indies. PLoS Negl Trop Dis. 2020 Jan 21;14(1):e0007940. doi: 10.1371/journal.pntd.0007940.

30. Ramos-Nino ME, Fitzpatrick DM, Eckstrom KM, Tighe S, Dragon JA, Cheetham S. The Kidney-Associated Microbiome of Wild-Caught Artibeus spp. in Grenada, West Indies. Animals (Basel). 2021 May 27;11(6):1571. doi: 10.3390/ani11061571.

31. Zamora-Vélez A, Cuadrado-Ríos S, Hernández-Pinsón A, Mantilla-Meluk H, Gómez-Marín JE. First Detection of Toxoplasma gondii DNA in a Wild Bat from Colombia. Acta Parasitol. 2020 Dec;65(4):969–973. doi: 10.2478/s11686-020-00222-1.

32. He B, Hu T, Yan X, Pa Y, Liu Y, Liu Y, Li N, Yu J, Zhang H, Liu Y, Chai J, Sun Y, Mi S, Liu Y, Yi L, Tu Z, Wang Y, Sun S, Feng Y, Zhang W, Zhao H, Duan B, Gong W, Zhang F, Tu C. Isolation, characterization, and circulation sphere of a filovirus in fruit bats. Proc Natl Acad Sci U S A. 2024 Feb 13;121(7):e2313789121. doi: 10.1073/pnas.2313789121.

33. Ge XY, Li JL, Yang XL, Chmura AA, Zhu G, Epstein JH, Mazet JK, Hu B, Zhang W, Peng C, Zhang YJ, Luo CM, Tan B, Wang N, Zhu Y, Crameri G, Zhang SY, Wang LF, Daszak P, Shi ZL. Isolation and characterization of a bat SARS-like coronavirus that uses the ACE2 receptor. Nature. 2013 Nov 28;503(7477):535–8. doi: 10.1038/nature12711.

34. Hu B, Zeng LP, Yang XL, Ge XY, Zhang W, Li B, Xie JZ, Shen XR, Zhang YZ, Wang N, Luo DS, Zheng XS, Wang MN, Daszak P, Wang LF, Cui J, Shi ZL. Discovery of a rich gene pool of bat SARS-related coronaviruses provides new insights into the origin of SARS coronavirus. PLoS Pathog. 2017 Nov 30;13(11):e1006698. doi: 10.1371/journal.ppat.1006698.

35. Zhou H, Ji J, Chen X, Bi Y, Li J, Wang Q, Hu T, Song H, Zhao R, Chen Y, Cui M, Zhang Y, Hughes AC, Holmes EC, Shi W. Identification of novel bat coronaviruses sheds light on the evolutionary origins of SARS-CoV-2 and related viruses. Cell. 2021 Aug 19;184(17):4380–4391.e14. doi: 10.1016/j.cell.2021.06.008.

36. Wang J, Pan YF, Yang LF, Yang WH, Lv K, Luo CM, Wang J, Kuang GP, Wu WC, Gou QY, Xin GY, Li B, Luo HL, Chen S, Shu YL, Guo D, Gao ZH, Liang G, Li J, Chen YQ, Holmes EC, Feng Y, Shi M. Individual bat virome analysis reveals co-infection and spillover among bats and virus zoonotic potential. Nat Commun. 2023 Jul 10;14(1):4079. doi: 10.1038/s41467-023-39835-1.

37. Aljofan M. Hendra and Nipah infection: emerging paramyxoviruses. Virus Res. 2013;177(2):119–126. doi:10.1016/j.virusres.2013.08.002

38. Léveillé AN, Bland SK, Carlton K, Larouche CB, Kenney DG, Brouwer ER, Lillie BN, Barta JR. Klossiella equi Infecting Kidneys of Ontario Horses: Life Cycle Features and Multilocus Sequence-Based Genotyping Confirm the Genus Klossiella Belongs In the Adeleorina (Apicomplexa: Coccidia). J Parasitol. 2019

39. Karunarathna SC, Dong Y, Karasaki S, Tibpromma S, Hyde KD, Lumyong S, Xu J, Sheng J, Mortimer PE. Discovery of novel fungal species and pathogens on bat carcasses in a cave in Yunnan Province, China. Emerg Microbes Infect. 2020 Dec;9(1):1554–1566. doi: 10.1080/22221751.2020.1785333.

40. Cai Y, Wang X, Zhang N, Li J, Gong P, He B, Zhang X. First report of the prevalence and genotype of Trypanosoma spp. in bats in Yunnan Province, Southwestern China. Acta Trop. 2019 Oct;198:105105. doi: 10.1016/j.actatropica.2019.105105.

41. Sawatsky, B., Grolla, A., Kuzenko, N., Weingartl, H., Czub, M., 2007. Inhibition of henipavirus infection by Nipah virus attachment glycoprotein occurs without cell-surface downregulation of ephrin-B2 or ephrin-B3. J Gen Virol 88 (Pt 2), 582–591.

42. Luby SP, Rahman M, Hossain MJ, Blum LS, Husain MM, Gurley E, Khan R, Ahmed BN, Rahman S, Nahar N, Kenah E, Comer JA, Ksiazek TG. Foodborne transmission of Nipah virus, Bangladesh. Emerg Infect Dis. 2006 Dec;12(12):1888–94. doi: 10.3201/eid1212.060732.

43. Luby SP, Hossain MJ, Gurley ES, Ahmed BN, Banu S, Khan SU, Homaira N, Rota PA, Rollin PE, Comer JA, Kenah E, Ksiazek TG, Rahman M. Recurrent zoonotic transmission of Nipah virus into humans, Bangladesh, 2001-2007. Emerg Infect Dis. 2009 Aug;15(8):1229–35. doi: 10.3201/eid1508.081237.

44. Li Y, Wang J, Hickey AC, et al. Antibodies to Nipah or Nipah-like viruses in bats, China. Emerg Infect Dis. 2008;14(12):1974–1976. doi:10.3201/eid1412.080359

45. Wu Z, Yang L, Yang F, et al. Novel Henipa-like virus, Mojiang Paramyxovirus, in rats, China, 2012. *Emerg Infect Dis*. 2014;20(6):1064-1066. doi:10.3201/eid2006.131022

46. Zhang XA, Li H, Jiang FC, et al. A Zoonotic Henipavirus in Febrile Patients in China. N Engl J Med. 2022;387(5):470–472. doi:10.1056/NEJMc2202705

47. Vashi NA, Reddy P, Wayne DB, Sabin B. Bat-associated leptospirosis. J Gen Intern Med. 2010 Feb;25(2):162-4. doi: 10.1007/s11606-009-1210-7.

48. Ballados-González GG, Sánchez-Montes S, Romero-Salas D, Colunga Salas P, Gutiérrez-Molina R, León-Paniagua L, Becker I, Méndez-Ojeda ML, Barrientos-Salcedo C, Serna-Lagunes R, Cruz-Romero A. Detection of pathogenic Leptospira species associated with phyllostomid bats (Mammalia: Chiroptera) from Veracruz, Mexico. Transbound Emerg Dis. 2018 Jun;65(3):773–781. doi: 10.1111/tbed.12802.

49. Bevans AI, Fitzpatrick DM, Stone DM, Butler BP, Smith MP, Cheetham S. Phylogenetic relationships and diversity of bat-associated Leptospira and the histopathological evaluation of these infections in bats from Grenada, West Indies. PLoS Negl Trop Dis. 2020 Jan 21;14(1):e0007940. doi: 10.1371/journal.pntd.0007940.

50. Ramos-Nino ME, Fitzpatrick DM, Eckstrom KM, Tighe S, Dragon JA, Cheetham S. The Kidney-Associated Microbiome of Wild-Caught Artibeus spp. in Grenada, West Indies. Animals (Basel). 2021 May 27;11(6):1571. doi: 10.3390/ani11061571.

51. Yang T, Yang W, Kuang G, Pan H, Han X, Yang L, Wang J, Feng Y. Prevalence and Characteristics of Novel Pathogenic Leptospira Species in Bats in Yunnan Province, China. Microorganisms. 2023 Jun 20;11(6):1619. doi: 10.3390/microorganisms11061619.

52. Lustermans JJM, Bjerg JJ, Schramm A, Marshall IPG. Phyllobacterium calauticae sp. nov. isolated from a microaerophilic veil transversed by cable bacteria in freshwater sediment. Antonie Van Leeuwenhoek. 2021;114(11):1877–1887. doi:10.1007/s10482-021-01647-y

53. Höhne K, Loose B, Seeliger HP. Isolation of Listeria monocytogenes in slaughter animals and bats of Togo (West Africa). Ann Microbiol (Paris). 1975 May-Jun;126A(4):501-7.

54. Povolyaeva O, Chalenko Y, Kalinin E, Kolbasova O, Pivova E, Kolbasov D, Yurkov S, Ermolaeva S. Listeria monocytogenes Infection of Bat Pipistrellus nathusii Epithelial cells Depends on the Invasion Factors InlA and InlB. Pathogens. 2020 Oct 22;9(11):867. doi: 10.3390/pathogens9110867.

55. Santana Lima VF, Rocha PA, Dias Silva MA, Beltrão-Mendes R, Ramos RAN, Giannelli A, Rinaldi L, Cringoli G, Estrela PC, Alves LC. Survey on helminths and protozoa of free-living Neotropical bats from Northeastern Brazil. Acta Trop. 2018 Sep;185:267–272. doi: 10.1016/j.actatropica.2018.06.002.

56. Jiang HH, Qin SY, Wang W, He B, Hu TS, Wu JM, Fan QS, Tu CC, Liu Q, Zhu XQ. Prevalence and genetic characterization of Toxoplasma gondii infection in bats in southern China. Vet Parasitol. 2014 Jul 14;203(3-4):318–21. doi: 10.1016/j.vetpar.2014.04.016.

57. Goodman RP, Ghabrial SA, Fichorova RN, Nibert ML. Trichomonasvirus: a new genus of protozoan viruses in the family Totiviridae. Arch Virol. 2011 Jan;156(1):171–9. doi: 10.1007/s00705-010-0832-8.

58. 58. ICTV (2024). The ICTV report on virus classification and taxon nomenclature. https://ictv.global/report.

59. Ivanova, N. V., Dewaard, J. R., & Hebert, P. D. N.. (2010). An inexpensive, automation-friendly protocol for recovering high-quality dna. Mol.ecol.notes, 6(4), 998–1002.DOI:10.1111/j.1471-8286.2006.01428.x.

60. Ratnasingham, S., & Hebert, P. D. (2007). bold: The Barcode of Life Data System (http://www.barcodinglife.org). Molecular ecology notes, 7(3), 355–364. 10.1111/j.1471-8286.2007.01678.x

61. Guindon S, Dufayard JF, Lefort V, Anisimova M, Hordijk W, Gascuel O. New algorithms and methods to estimate maximum-likelihood phylogenies: assessing the performance of PhyML 3.0. Syst Biol. 2010 May;59(3):307–21. doi: 10.1093/sysbio/syq010.

62. Fu L, Niu B, Zhu Z, Wu S, Li W. CD-HIT: accelerated for clustering the next-generation sequencing data. Bioinformatics. 2012 Dec 1;28(23):3150–2. doi: 10.1093/bioinformatics/bts565.

63. Langmead B, Salzberg SL. Fast gapped-read alignment with Bowtie 2. Nat Methods. 2012 Mar 4;9(4):357–9. doi: 10.1038/nmeth.1923.

64. Buchfink B, Xie C, Huson DH. Fast and sensitive protein alignment using DIAMOND. Nat Methods. 2015 Jan;12(1):59–60. doi: 10.1038/nmeth.3176.

65. Li D, Liu CM, Luo R, Sadakane K, Lam TW. MEGAHIT: an ultra-fast single-node solution for large and complex metagenomics assembly via succinct de Bruijn graph. Bioinformatics. 2015 May 15;31(10):1674–6. doi: 10.1093/bioinformatics/btv033.

66. Clewley JP. Macintosh sequence analysis software. DNAStar’s LaserGene. Mol Biotechnol. 1995 Jun;3(3):221–4. doi: 10.1007/BF02789332.

67. Li H. A statistical framework for SNP calling, mutation discovery, association mapping and population genetical parameter estimation from sequencing data. Bioinformatics. 2011 Nov 1;27(21):2987–93. doi: 10.1093/bioinformatics/btr509.

68. Kearse M, Moir R, Wilson A, Stones-Havas S, Cheung M, Sturrock S, Buxton S, Cooper A, Markowitz S, Duran C, Thierer T, Ashton B, Meintjes P, Drummond A. Geneious Basic: an integrated and extendable desktop software platform for the organization and analysis of sequence data. Bioinformatics. 2012 Jun 15;28(12):1647–9. doi: 10.1093/bioinformatics/bts199.

69. Segata N, Waldron L, Ballarini A, Narasimhan V, Jousson O, Huttenhower C. Metagenomic microbial community profiling using unique clade-specific marker genes. Nat Methods. 2012 Jun 10;9(8):811–4. doi: 10.1038/nmeth.2066.

70. Katoh K, Standley DM. MAFFT multiple sequence alignment software version 7: improvements in performance and usability. Mol Biol Evol. 2013 Apr;30(4):772–80. doi: 10.1093/molbev/mst010.

71. Capella-Gutiérrez S, Silla-Martínez JM, Gabaldón T. trimAl: a tool for automated alignment trimming in large-scale phylogenetic analyses. Bioinformatics. 2009 Aug 1;25(15):1972–3. doi: 10.1093/bioinformatics/btp348

